# Large-scale differential gene expression analysis identifies genes associated with Bipolar Disorder in post-mortem brain

**DOI:** 10.1101/770529

**Authors:** Mohamed N. Omar, Mohamed Youssef, Mohamed Abdellatif

## Abstract

**Background and purpose:** Bipolar disorder (BD) is a common psychiatric disorder with high morbidity and mortality. Several polymorphisms have been found to be implicated in the pathogenesis of BD, however, these loci have small effect sizes that fail to explain the high heritability of the disease. Here, we provide more insights into the genetic basis of BD by identifying the differentially expressed genes (DEGs) and their associated pathways and biological processes in post-mortem brain tissues of patients with BD.

**Methods:** Eight datasets were eligible for the differential expression analysis. We used six datasets for the discovery of the gene signature and used the other two for independent validation. We performed the multi-cohort analysis by a random-effect model using R and MetaIntegrator package.

**Results:** The initial analysis resulted in the identification of 126 DEGs (30 up-regulated and 94 down-regulated). We refined this initial signature by a forward search process and resulted in the identification of 22 DEGs (6 up-regulated and 16 down-regulated). We validated the final gene signature in the independent datasets and resulted in an Area Under the ROC Curve (AUC) of 0.756 and 0.76, respectively. We performed gene set enrichment analysis (GSEA) which identified several biological processes and pathways related to BD including Ca transport, inflammation and DNA damage response.

**Conclusion:** Our findings support the previous findings that link BD pathogenesis to abnormalities in glial inflammation and calcium transport and also identify several other biological processes not previously reported to be associated with the development of the disease. Such findings will improve our understanding of the genetic basis underlying BD and may have future clinical implications.

## INTRODUCTION

Bipolar disorder (BD) is a common psychiatric disorder characterized by alternating periods of depression and abnormally elevated mood associated with cognitive impairments and increased impulsivity [1, 2, 3]. The lifetime prevalence of Bipolar disorder is approximately 2.4% with average age of onset of 25 years [4].

Aside from mental and cognitive impairments, BD is also associated with high cardiovascular morbidity and mortality [5, 6, 7]. This association was explained by the co-occurrence of risk factors or due to side effects of psychotropic medications [8].

On the neuro-anatomical level, neuro-imaging studies link BD to abnormalities in various areas of the brain, especially the frontal, prefrontal and parietal cortices with cortical thinning and increased ventricular volume [9, 10].

On the genetic level, twin studies showed that BD is a highly heritable disease with heritability reaching up to 80% [11, 12]. The risk of getting the disease is 10-fold higher in the first degree relatives of those affected with BD compared to normal population. Although recent studies showed that several susceptibility loci with small effect sizes are associated with BD [13, 14], there is no much evidence supporting the role of specific genes in the pathogenesis of the disease [15].

The molecular mechanisms underlying the etiology of bipolar disorder remain unclear with recent evidence pointing toward a significant role of immune disturbances [16, 17], disrupted Ca dynamics [18, 19] and oxidative DNA damage in the pathogenesis of BD [20].

In this study, we attempt to identify an accurate and consistent gene signature of BD in post-mortem brain rather than detecting polymorphisms or copy number variants (CNVs). We also attempt to identify the biological pathways and processes of this gene signature. For this purpose, we analyze microarray gene expression data to identify the deferentially expressed genes (DEGs) between BD and healthy controls. Detecting those genes and pathways would offer a better understanding of the molecular pathogenesis of this disease and may have future therapeutic implications by targeting these genes or pathways.

## METHODS

### Data Collection

We performed an extensive search of the Gene Expression Omnibus (GEO) database [21] looking for microarray gene expression datasets from patients with BD and from healthy controls. We used the following search term “Bipolar disorder” and the following filters: entry type (series), study type (Expression profiling by array) and organism (Homo sapiens). This resulted in a total of 39 studies which were then filtered to include only gene expression studies in post-mortem brain tissues only (frontal, prefrontal and parietal cortices).

A total of eight datasets were eligible for the downstream analysis (Table 1). In all datasets, we removed samples from patients with schizophrenia and major depressive disorder and kept only samples from BD patients and controls. This resulted in a total of 509 samples with 248 BD and 261 control samples. We used six datasets (378 samples, 74.3% of the total samples) for discovery of the BD gene signature and kept two datasets (131 samples, 25.7% of the total samples) out for independent validation of the signature performance.

**Table 1:** Collected data sets showing the number of samples with the number of BD cases.

| GEO accession | Type of experiment | Platform | Number of samples (BD cases) | Tissue source |
| --- | --- | --- | --- | --- |
| **Discovery** |  |  |  |  |
| **GSE120340** [45] | Expression profiling by array | GPL16311 | 20 (10) | Dorsolateral pre-frontal cortex |
| **GSE87610** [46] | Expression profiling by array | GPL13667 | 145 (73) | Dorsolateral pre-frontal cortex |
| **GSE12654** [47] | Expression profiling by array | GPL8300 | 26 (11) | Anterior pre-frontal cortex |
| **GSE12649** [47] | Expression profiling by array | GPL96 | 67 (33) | Dorsolateral pre-frontal cortex |
| **GSE62191** [48] | Expression profiling by array | GPL4133 | 59 (29) | Frontal cortex |
| **GSE5388** [49] | Expression profiling by array | GPL96 | 61 (30) | Dorsolateral pre-frontal cortex |
| **Validation** |  |  |  |  |
| **GSE35977** [50, 51] | Expression profiling by array | GPL6244 | 95 (45) | Parietal cortex |
| **GSE53987** [52] | Expression profiling by array | GPL570 | 36 (17) | Dorsolateral pre-frontal cortex |

### The Meta-Analysis

We used R Programming language [22] and the “MetaIntegrator” package [23] which utilizes a gene expression meta-analysis workflow described by Haynes et al [24]. The package can be used to retrieve the datasets from GEO, then it automatically checks if these datasets are appropriately normalized and log2-scaled. We found that two datasets (GSE12654 and GSE12649) were initially normalized using median centering so we downloaded the raw data and normalized them using Robust Multiarray Average (RMA) method which is the same method used to normalize the other Affymetrix datasets. All datasets were on log2-scale before conducting the meta-analysis.

The MetaIntegrator approach computes a Hedges effect size [25]for each gene in each dataset. These effect sizes are then pooled across all the datasets (Supp. figures1,2) using a random-effect model by assuming that results from each study is drawn from a single distribution and that each inter-study difference is just a random effect. The approach computes the log sum of p-values that each gene is up/down-regulated, then combine p-values using Fisher’s method and finally performs Benjamini-Hochberg false discovery rate (FDR) correction across all genes [26].

In our analysis, a gene would be deferentially expressed if the absolute value of its effect size is greater than zero, the FDR is less than 5% across all discovery datasets and it has to be significantly up/down regulated in all of the six training datasets with a heterogeneity P-value cutoff of 0.05 [27]. All discovery steps were conducted on the training datasets only.

In order to optimize the initial gene signature, we performed a Forward Search process by taking the initial gene set, adding one gene at a time and calculating the weighted Area Under the ROC curve (AUC) resulting from the addition of this gene. Weighted AUC is calculated as:

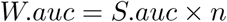

where W.auc is the weighted AUC, S.auc is the sum of AUC of each data set and n is the number of samples in the data set.

This process is repeated for each gene until the stopping threshold (0 in our case) is reached and the final set of genes will be those who contributed the most to the weighted AUC.

### Validation of the Resulting Gene Signature

We tested the performance and consistency of the resulting BD gene signature in the two independent cohorts (Table 1). We used the Area Under the Receiver Operating Characteristic Curve (AUC), Area Under the Precision Recall Curve (AUPRC) and Matthews correlation coefficient (MCC) as evaluation metrics.

### Enrichment Analysis

Functional Enrichment analysis was performed using Enrichr R package [28, 29] and the following databases: “GO-Biological-Process-2018” and “KEGG-2019-Human”. The goal was to discover the biological processes and pathways enriched in the discovered gene set.

## RESULTS

### The BD Gene Signature

The initial meta-analysis resulted in the identification of 124 DEGs (30 up-regulated and 94 down-regulated). We refined this initial signature by using a forward search process which resulted in the identification of 22 DEGs (6 up-regulated and 16 down-regulated). Figure (1) shows the summary AUC of the discovery datasets with the initial gene signature and the summary AUC with the final signature resulting from forward search.

**Figure 1:**
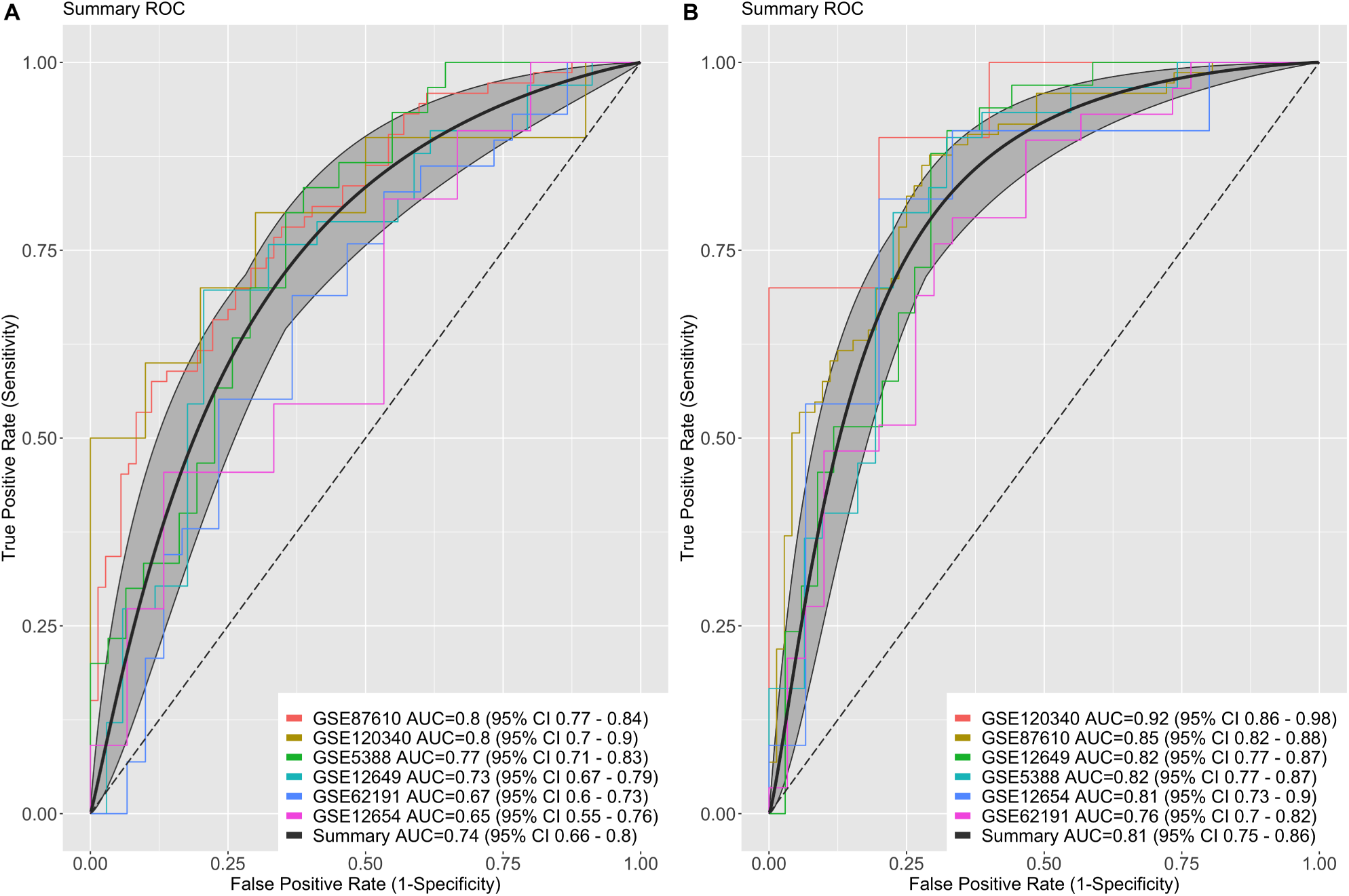
Summary AUC in the training datasets. with the initial gene signature (A) and the final gene signature resulting from forward search (B)

Of the 22 DEGs, 6 genes were up-regulated (*CTBP2, ELK1, ARHGAP5, RNASE4, MAOA, GOLGA8A*) (Supp. figure1) and 16 were down-regulated (*SST, IGFBP6, CD53, RABIF, DDX18, NXF1, CXCL12, ETV5, HCLS1, NFYC, SULF1, ZFYVE9, CSE1L, DNAJC7, FDFT1, TSN*) (Table 2) (Supp. figure2).

**Table 2:** The BD signature genes. 13 up-regulated and 9 down-regulated genes.

| Gene Symbol | Effect size | Effect size SE | Effect size p-value | Effect size FDR |
| --- | --- | --- | --- | --- |
| **Up-regulated genes** |  |  |  |  |
| *CTBP2* | 0.295 | 0.046 | 1.08e-10 | 7.56e-07 |
| *ARHGAP5* | 0.308 | 0.056 | 5.13e-08 | 9.82e-05 |
| *GOLGA8A* | 0.192 | 0.044 | 1.58e-05 | 0.004 |
| *RNASE4* | 0.321 | 0.084 | 0.0001 | 0.012 |
| *MAOA* | 0.339 | 0.102 | 0.0009 | 0.037 |
| *ELK1* | 0.327 | 0.099 | 0.0009 | 0.037 |
| **Down-regulated genes** |  |  |  |  |
| *NFYC* | -0.341 | 0.069 | 6.65e-07 | 0.0005 |
| *NXF1* | -0.385 | 0.089 | 1.37e-05 | 0.0043 |
| *SST* | -0.652 | 0.151 | 1.54e-05 | 0.0043 |
| *DDX18* | -0.258 | 0.06 | 1.53e-05 | 0.0043 |
| *ETV5* | -0.430 | 0.105 | 4.22e-05 | 0.0071 |
| *FDFT1* | -0.234 | 0.057 | 4.42e-05 | 0.0073 |
| *ZFYVE9* | -0.279 | 0.069 | 5.11e-05 | 0.0076 |
| *TSN* | -0.266 | 0.066 | 4.99e-05 | 0.0076 |
| *IGFBP6* | -0.419 | 0.104 | 5.9e-05 | 0.0082 |
| *CSE1L* | -0.247 | 0.062 | 7.25e-05 | 0.0087 |
| *SULF1* | -0.382 | 0.098 | 9.69e-05 | 0.0104 |
| *CXCL12* | -0.271 | 0.070 | 0.0001 | 0.011 |
| *CD53* | -0.442 | 0.115 | 0.0001 | 0.0117 |
| *RABIF* | -0.441 | 0.117 | 0.0002 | 0.014 |
| *HCLS1* | -0.267 | 0.078 | 0.0006 | 0.03 |
| *DNAJC7* | -0.313 | 0.098 | 0.0014 | 0.047 |

| Gene Symbol | Effect size | Effect size SE | Effect size p-value | Effect size FDR |
| --- | --- | --- | --- | --- |

### Strong Performance in the Two Validation Datasets

We measured the discriminatory performance of our gene signature by testing it in two independent datasets (N=131) (Fig. 2). In the first validation dataset (GSE35977), the signature achieved an AUC of 0.756 (95%CI: 0.66-0.85) (Fig. 2), AUPRC of 0.70 (95%CI: 0.57-0.83), accuracy of 73% (95% CI: 63%-81%, P=5.3e-05), sensitivity of 76% and MCC of 0.46.

**Figure 2:**
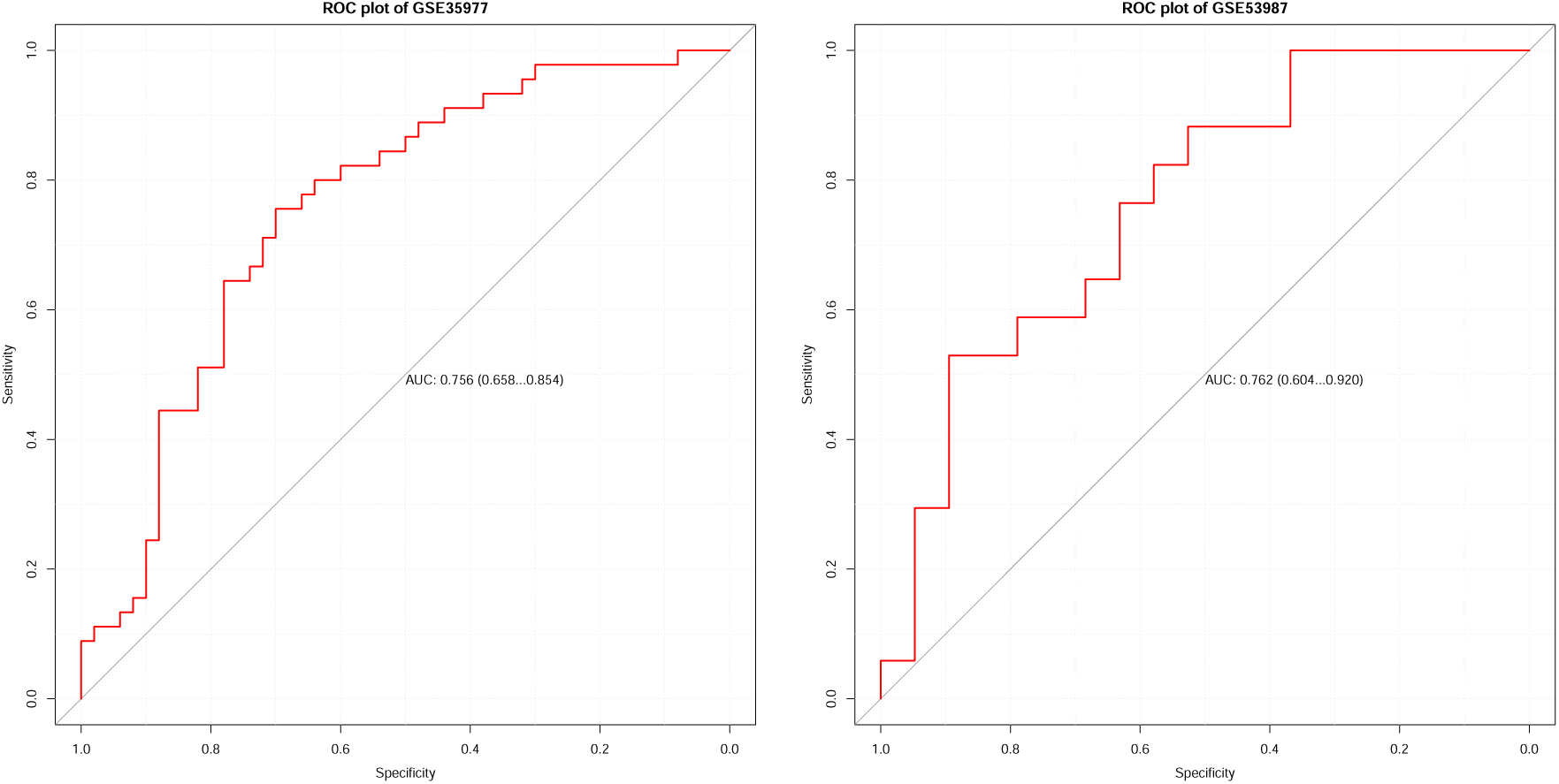
ROC curves of the two validation data sets. Strong classification performance (AUC = 0.756 and 0.76 respectively).

In the second validation dataset (GSE53987), the signature achieved an AUC of 0.762 (95%CI: 0.60-0.92) (Fig. 3), AUPRC of 0.704 (95%CI: 0.49-0.72), accuracy of 72% (95%CI: 55%-86%, P=0.014), sensitivity of 53% and MCC of 0.46.

**Figure 3:**
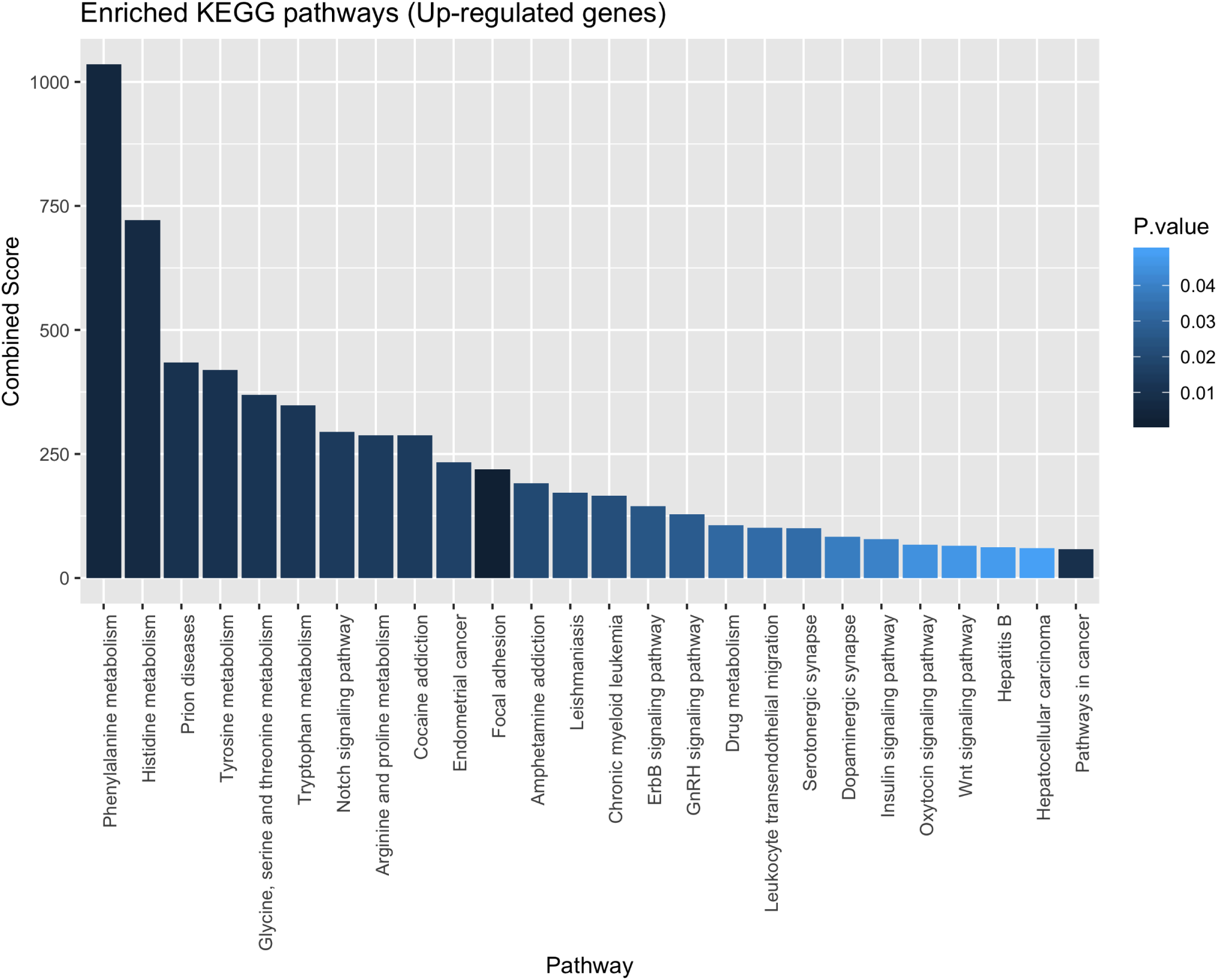
KEGG pathways of the 6 up-regulated genes.

The signature performance in the validation datasets showed great consistency with the performance in the discovery datasets with minimal overfitting (Summary AUC = 0.81, 95% CI:0.75-0.86).

### Enrichment Analysis

Gene set enrichment analysis (GSEA) has revealed several significant (P*<* 0.05) KEGG pathways and GO biological processes enriched in the discovered gene signature. The six up-regulated genes were associated with 26 KEGG pathways (Fig. 3) (Supp.table1) and 18 GO biological processes (Fig. 5) (Supp. table2). The 16 down-regulated genes were associated with 3 KEGG pathways (Fig. 4) (Supp. table3) together with 102 GO biological processes (Fig. 6) (Supp. table4).

**Figure 4:**
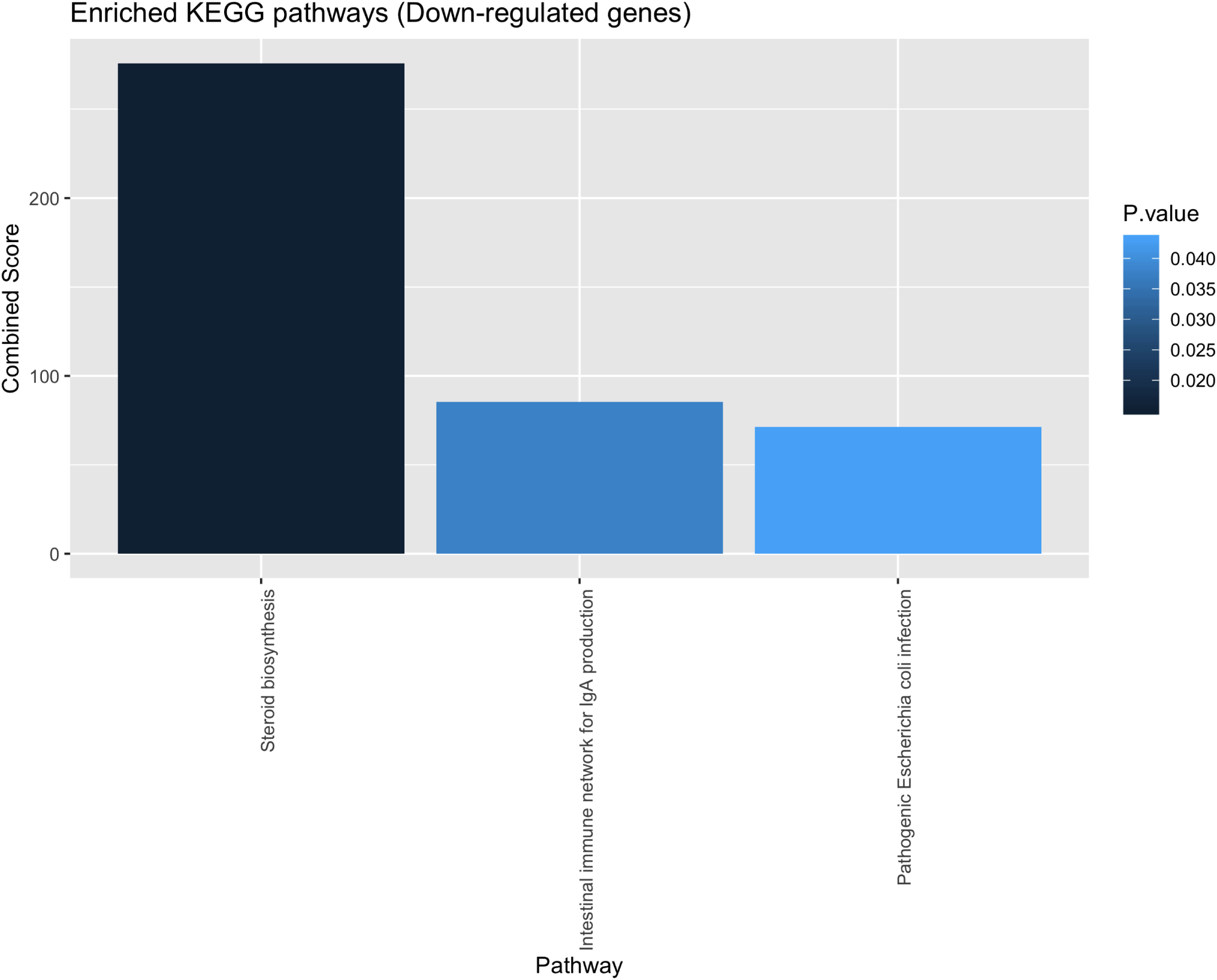
KEGG pathways of the 16 down-regulated genes.

**Figure 5:**
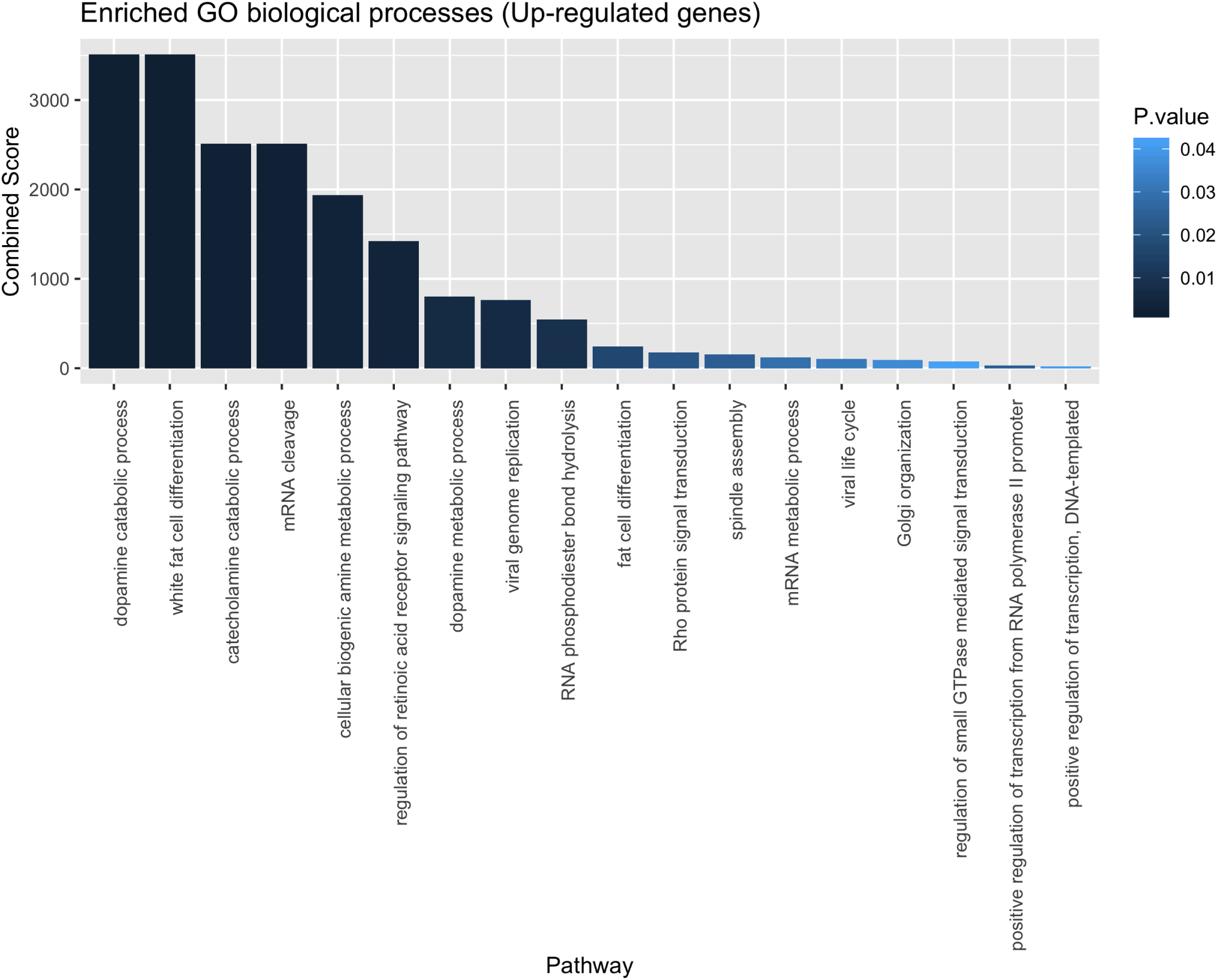
GO biological processes of the 6 up-regulated genes.

**Figure 6:**
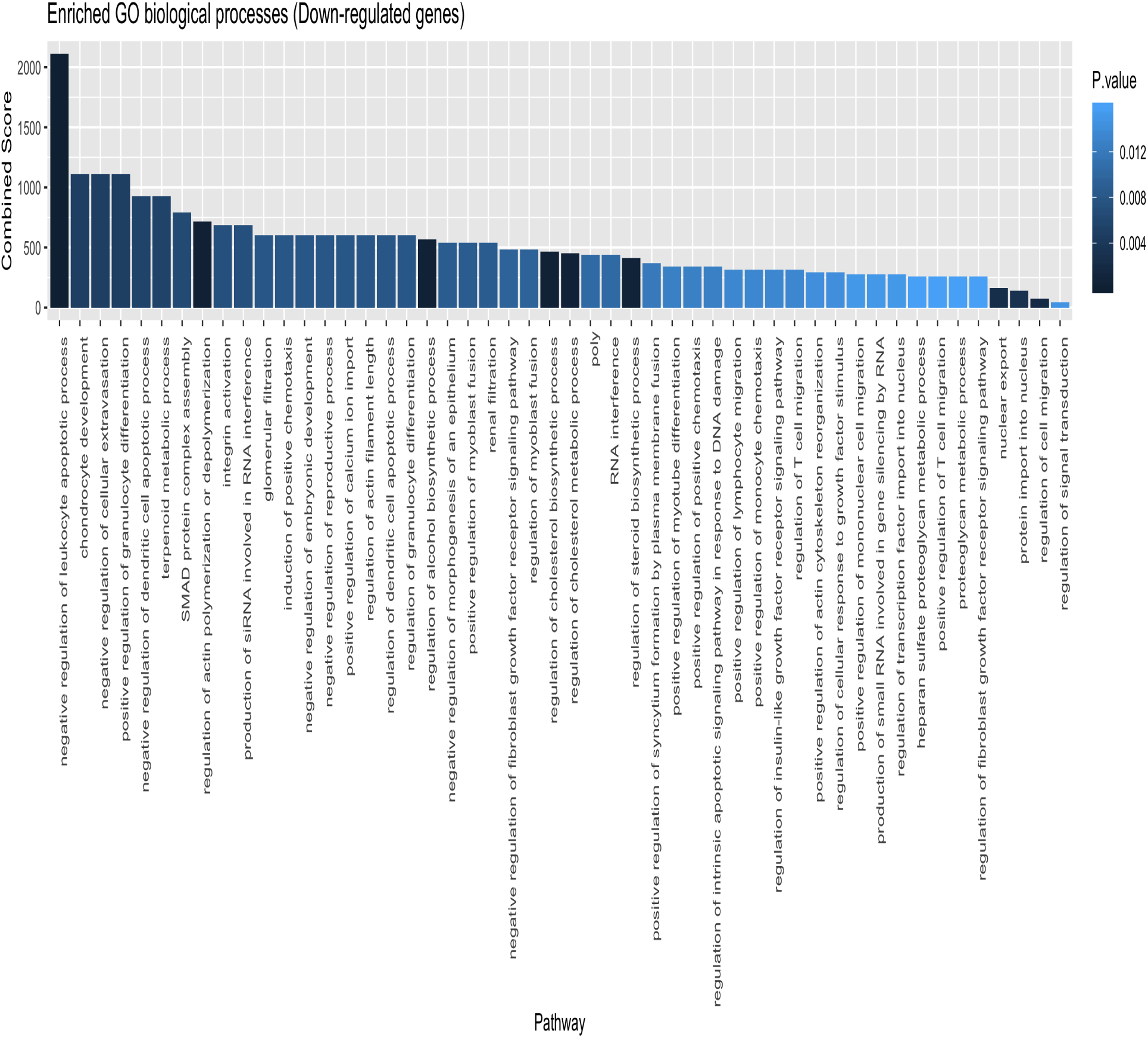
GO biological processes of the 16 down-regulated genes.

## DISCUSSION

Bipolar disorder is a highly heterogeneous disease and the genetic basis underlying its pathophysiology is complex [30]. Several genome-wide association studies (GWAS) have identified several copy number variants (CNVs) and single-nucleotide polymorphisms (SNPs) that seem to play a role in the high heritability of the disease but each of these variants has a small effect [31, 32]. Moreover, most of these studies failed to identify specific genes that may have a large effect in the pathogenesis of the disease, most likely due to small sample size.

Here we attempted to overcome the problem of inadequate sample size by integrating multiple microarray gene expression studies in a meta-analysis process to achieve high statistical power. Through this large-scale gene expression analysis of 509 post-mortem brain samples, we managed to identify six up-regulated and sixteen down-regulated genes in the post-mortem brain tissues of patients with BD. The resulting gene signature proved to be accurate and consistent when tested in two independent validation datasets.

GSEA identified important pathways and processes enriched in the discovered BD gene signature. The up-regulated genes were particularly associated with important cell signaling pathways like notch and Wnt signaling pathways which were previously reported to have a significant role in the pathogenesis of BD together with other psychiatric disorders [33, 34, 35, 36]. Additionally they were also associated with pathways involved in inflammatory cell motility like focal adhesion and leukocyte transendothelial migration pathways. The GO biological processes enriched in the up-regulated genes included dopamine and catecholamine catabolic processes together with processes that regulate signal transduction like Rho protein signal transduction and GTPase-mediated signal transduction supporting the current evidence that links BD to alterations in the cytoskeletal dynamics [37, 38]. Other biological processes included those regulating transcription and mRNA cleavage.

On the other hand, the down-regulated genes were associated with processes that regulate apoptosis, Ca transport, cell migration, granulocytes differentiation, chemotaxis and lymphocytes migration, supporting the notion that glial inflammation/loss, imbalances in the apoptotic process and Ca homeostasis may have a role in the pathogenesis of BD [39, 40, 41, 42, 18].

Most of these biological processes and pathways enriched in the BD gene signature overlap with those responsible for the development of atherosclerosis [43, 44]. This suggests that the association between BD and cardiovascular diseases may have some genetic basis rather than shared environmental risk factors or side effects of psychotropic medications.

One limitation of this study is the lack of experimental validation in another independent cohort. Although our signature showed great performance and consistency in the validation datasets, this assessment was purely computational and it needs experimental validation using real-time PCR to assess its clinical validity.

## CONCLUSION

In this study, we provide more insights into the genetic basis of BD through the identification of DEGs in post-mortem brain tissues of BD patients compared with normal controls. Our discovered BD gene signature proved to be accurate and consistent as shown by its performance in the independent validation cohorts. The BD signature genes were associated with a number of biological processes and pathways, most importantly the pathways regulating Ca transport, immune response, oxidative damage and inflammatory processes. These pathways are also involved in the pathogenesis of atherosclerosis, cardiovascular diseases and metabolic syndrome. Such findings, together with previous research will provide a more clear understanding of the molecular pathophysiology of BD and may have future therapeutic potentials by targeting these specific genes and pathways.

## List of Abbreviations

BD: Bipolar Disorder.
DEGs: Differentially Expressed Genes.
GEO: Gene Expression Omnibus.
ROC: Receiver Operating Characteristic.
AUC: Area Under the ROC Curve.
AUPRC: Area Under the Precision Recall Curve.
MCC: Matthews Correlation Coefficient.
GO: Gene Ontology.
KEGG: Kyoto Encyclopedia of Genes and Genomes.

## Ethics approval and consent to participate

Not applicable

## Consent for publication

Not applicable

## Availability of data and materials

All data sets used in this study are publicly available on the Gene Expression Omnibus (GEO) data base under the corresponding accession number. The code for this analysis is available on GitHub and can accessed using the following link: https://github.com/MohamedOmar2020/BD

## Author’s contributions

MO conceived the research question and performed the analysis. MY and MA collected the data sets. All authors contributed equally to the manuscript writing, read and approved the final manuscript.

## Acknowledgements

We thank Dr.Mohamed Tarek Badr (University Medical Center Freiburg, Germany) for his valuable suggestions that greatly improved the manuscript.

## Additional Files

**Supp. figure1— EffectSizes-UpRegulated**

PDF file showing forest plots of the effect sizes of each of the 6 up-regulated genes.

**Supp. figure2— EffectSizes-DownRegulated**

PDF file showing forest plots of the effect sizes of each of the 16 down-regulated genes.

**Supp. table1 — KEGG-UpRegulated**

CSV table showing the significant KEGG pathways of the 6 up-regulated genes.

**Supp. table2— GO-UpRegulated**

CSV table showing the significant GO biological processes of the 6 up-regulated genes.

**Supp. table3 — KEGG-DownRegulated**

CSVtable showing the significant KEGG pathways of the 16 down-regulated genes.

**Supp. table4 — GO-DownRegulated**

CSVtable showing the significant GO biological processes of the 16 down-regulated genes.

