## Supplementary figures and images for "Large-scale differential gene expression analysis identifies genes associated with Bipolar Disorder in post-mortem brain"

### Supp. figure1

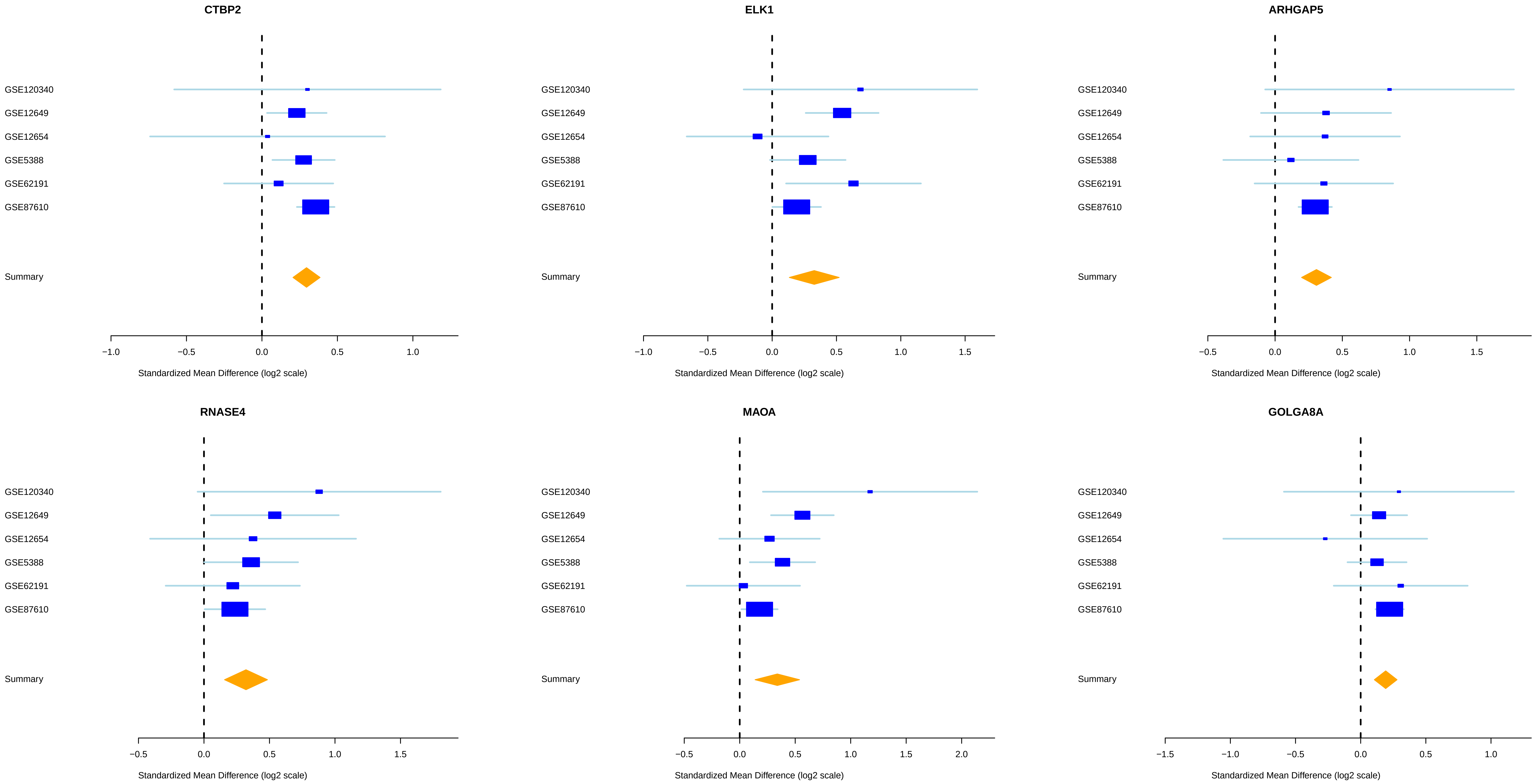

### Supp. figure2

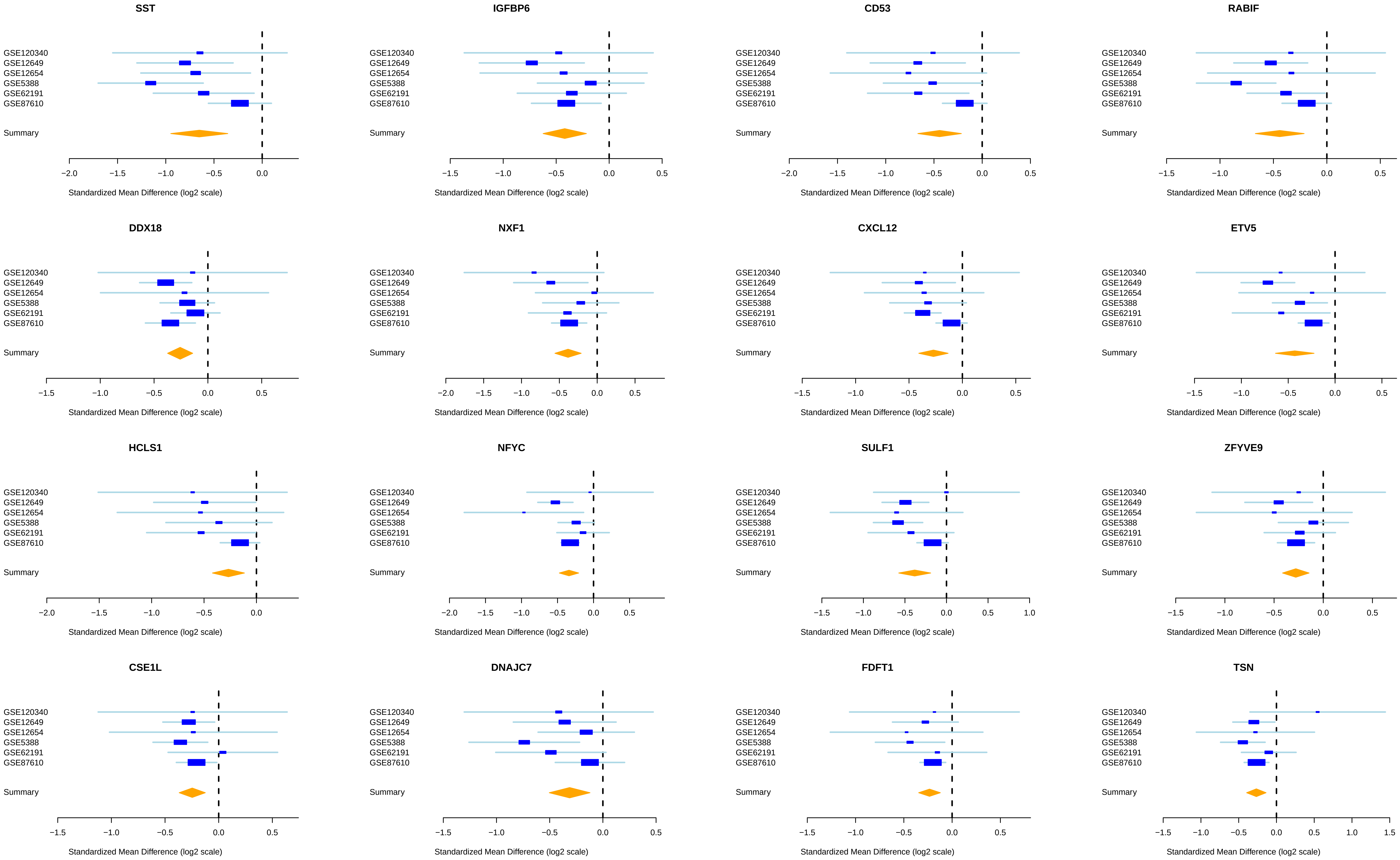
